# Stoichiometric constraints modulate the effects of temperature and nutrients on biomass distribution and community stability

**DOI:** 10.1101/589895

**Authors:** Arnaud Sentis, Bart Haegeman, José M. Montoya

## Abstract

Temperature and nutrients are two of the most important drivers of global change. Both can modify the elemental composition (i.e. stoichiometry) of primary producers and consumers. Yet their combined effect on the stoichiometry, dynamics, and stability of ecological communities remains largely unexplored. To fill this gap, we extended the Rosenzweig-MacArthur consumer-resource model by including thermal dependencies, nutrient dynamics, and stoichiometric constraints on both the primary producer and the consumer. We found that stoichiometric constraints dampen the paradox of enrichment and increased persistence at high nutrient levels. Nevertheless, they also reduced consumer persistence at extreme temperatures. Finally, we also found that stoichiometric constraints can strongly influence biomass distribution across trophic levels by modulating consumer assimilation efficiency and resource growth rates along the environmental gradients. In the Rosenzweig-MacArthur model, consumer biomass exceeded resource biomass for most parameter values whereas, in the stoichiometric model, consumer biomass was strongly reduced and sometimes lower than resource biomass. Our findings highlight the importance of accounting for stoichiometric constraints as they can mediate the temperature and nutrient impact on the dynamics and functioning of ecological communities.

## Introduction

Temperature and nutrients regulate many biological processes, including species geographical distribution, primary production, species interactions, and energy and material fluxes (Falkowski et al. 1998; Enquist et al. 1999; Elser et al. 2007; Thomas et al. 2017). They are at the core of several ecological theories. While temperature is a fundamental component of metabolic scaling theory (Brown et al. 2004), nutrients are at the core of resource competition theory (Tilman 1982) and ecological stoichiometry (i.e. the element composition of organisms) theory (Sterner & Elser 2002). Cross et al. (2015) suggested that a better understanding of the interactions between temperature and nutrients is crucial for developing realistic predictions about ecological responses to multiple drivers of global change, including climate warming and elevated nutrient supply. Nutrients can modulate the effects of warming on communities directly by altering primary production, and/or indirectly by changing the elemental composition of primary producers. Conversely, thermal effects on trophic interaction strengths (i.e. the per capita effect of predators on prey population densities) and on consumer energetic efficiencies (i.e. ingestion relative to metabolic demand) depend on both the quantity and quality of their resources. While Cross et al. (2015) provided a road map on how to investigate the combined effects of temperature and nutrient on ecological processes, we still lack an integrative theory to better understand how the links between stoichiometry, nutrient enrichment, and temperature influence the dynamics and stability of multispecies communities. Such a theory will allow us to understand how and when stoichiometric variation modulates the consequences of single and combined components of global change on trophic interactions, community dynamics, and ecosystem functioning.

Predicting the effects of global warming and nutrient changes on ecosystems is challenging as species are embedded within communities of multiple interacting species (Petchey et al. 1999; Tylianakis et al. 2008; Montoya & Raffaelli 2010; Gilbert et al. 2014). Increased resource availability (hereafter: enrichment) and warming can jointly affect food-web stability and structure by modifying the strength of trophic interactions (O’Connor et al. 2009; Binzer et al. 2012; Kratina et al. 2012; Sentis et al. 2014; Binzer et al. 2016). Enrichment typically increases energy flux from resources to higher trophic levels which often leads to the well-known paradox of enrichment where the amplitude of population fluctuations increase with nutrients, leading to extinctions at high nutrient concentrations (Rosenzweig 1971; Rip & McCann 2011; Gilbert et al. 2014). Nevertheless, most consumer species become less efficient at processing matter and energy at warmer temperatures as their metabolic rates often increase faster with temperature than their feeding rates (Vucic-Pestic et al. 2011; Fussmann et al. 2014; Iles 2014). This reduction of energetic efficiency lessens energy flow between trophic levels and hence stabilizes food-web dynamics by reducing population fluctuations (Rip & McCann 2011; Binzer et al. 2012; Gilbert et al. 2014). As a result, mild warming may alleviate the paradox of enrichment by decreasing consumer energetic efficiency (Binzer et al. 2012; Sentis et al. 2017).

The theoretical expectations and results described above have already improved our ability to understand and predict the effects of temperature and enrichment on food webs (Boit et al. 2012; Tabi et al. 2019). However, most previous studies using metabolic scaling theory assumed that nutrient enrichment lead to an increase in resource carrying capacity without influencing resource elemental composition (Vasseur & McCann 2005; Binzer et al. 2012; Gilbert et al. 2014; Binzer et al. 2016; Sentis et al. 2017). Yet nutrient enrichment effects are more complex. The elemental composition of primary producers is likely to be altered, in response to the supplies of energy and materials relative to their growth and nutrient intake rates (Rastetter et al. 1997; Robert W. Sterner et al. 1997; Finkel et al. 2009). This, in turn, can affect the dynamics of the producer population and the herbivores feeding on it. For instance, previous modelling studies showed that introducing stoichiometric heterogeneity in predator-prey population dynamic models can dampen the negative effect of nutrient enrichment on system persistence by reducing population biomass fluctuations (Andersen 1997; Loladze et al. 2000; Andersen et al. 2004; Elser et al. 2012). More generally, the stoichiometric flexibility of primary producers, in particular the flexibility in carbon to nutrient ratios (e.g. C:N or C:P), has important implications for animal feeding behaviour (White 1993), consumer population stability (White 1993; Sterner & Hessen 1994; Hessen et al. 2002), community structure (Andersen 1997), and ecosystem processes such as biogeochemical cycling (Andersen 1997; Hessen et al. 2004).

Previous theoretical and empirical studies reported that stoichiometric variations can have a strong influence on the stability of consumer-resource interactions (Andersen 1997; Andersen et al. 2004; Diehl et al. 2005; Elser et al. 2012). For instance, populations of crustacean Daphnia feeding on low quality (i.e. low nutrient: carbon ratio) algae cannot persist even when resource quantity is not a limiting factor (Elser et al. 2007). Consumer extinction is explained by the fact that the consumer assimilation efficiency is, for most organisms, a function of resource quality (Elser et al. 2000). When resource quality is low, the consumers assimilate only few nutrients relative to the biomass they ingest, which limits their growth and reproduction (Elser et al. 2000; Elser et al. 2012). Temporal variations in resource quality can stabilize the system by weakening interaction strength and dampening population fluctuations (Andersen et al. 2004; Diehl et al. 2005)but see(Loladze et al. 2000; Elser et al. 2012). However, it remains unclear whether and how temporal variations in the elemental composition of primary producers and consumers can modulate the effects of temperature and nutrients on important community features such as stability and biomass distribution across trophic levels. Previous studies indicated that the spatial and temporal intraspecific variations in the elemental composition of primary producers are expected to increase in response to global change drivers such as temperature, CO2, and nutrient availability (Bezemer & Jones 1998; Woods et al. 2003; Finkel et al. 2009). This increased variation can be of importance for both primary producer and consumer populations as the growth rate of primary producers is well known to depend on their elemental composition (Droop 1974) as is the assimilation efficiency of the consumers (Sterner & Elser 2002).

Altogether, previous studies indicated that both temperature and stoichiometric variations can have important effects on species interactions and community dynamics (Andersen et al. 2004; Diehl et al. 2005; Fussmann et al. 2014; Binzer et al. 2016; Sentis et al. 2017). However, the effects of temperature and nutrient stoichiometry on food web dynamics and stability have only been studied in isolation. Recent theory by Uszko et al. (2017) showed that considering nutrient dynamics can help to better understand the influence of temperature on consumer-resource population dynamics and resource carrying capacity. Nevertheless, they considered that the elemental composition of both the resource and the consumer are constant and independent of temperature and nutrient dynamics. This contrasts with the empirical observation that resource elemental composition is flexible and can vary with both temperature and nutrient dynamics (Droop 1974; Elser et al. 2000; Woods et al. 2003). Here we thus focused on the combined effects of temperature and nutrients on the stoichiometry of primary producers and how this affects community stability and biomass distribution across trophic levels in a consumer-resource system. Understanding the determinants of stability and biomass distribution has been at the core of ecology for a long time (Elton (1927), Lindeman (1942)). Recent theory aims at explaining empirical observations of trophic pyramids (i.e. population biomass decreases with trophic levels), inverted trophic pyramids (i.e. population biomass increases with trophic levels), trophic cascades and the link between biomass distribution and stability (McCauley et al. 2018; Barbier & Loreau 2019).

Here, we used the Rosenzweig-MacArthur model as a baseline non-stoichiometric model because this model is one of the most studied models used to investigate the effects of temperature and nutrient enrichment on community dynamics (Vasseur & McCann 2005; Binzer et al. 2012; Fussmann et al. 2014; Sentis et al. 2017). Inspired by previous temperature-independent stoichiometric consumer-resource models (Andersen 1997; Andersen et al. 2004; Diehl et al. 2005), we then extended the Rosenzweig-MacArthur model to account for nutrient dynamics as well as for the simultaneous dependence of community dynamics on temperature and flexible resource stoichiometry. Our objective here was not to develop a complex and very realistic stoichiometric model that would include additional important abiotic and biotic features such as light intensity (Diehl 2007) or compensatory feeding (Cruz-Rivera & Hay 2000). Instead, we aimed at introducing two fundamental stoichiometric features (i.e. stoichiometric flexibility and stoichiometric imbalance) and investigate how these stoichiometric considerations can change predictions of the Rosenzweig-MacArthur model. We thus used our extended Rosenzweig-MacArthur model to predict the effects of warming and nutrient enrichment on population dynamics and biomass distribution across trophic levels and compared these predictions with the predictions of the nonstoichiometric Rosenzweig-MacArthur model. We particularly addressed two questions: (i) How do stoichiometric constraints modulate the effects of enrichment and warming on community stability and persistence? and (ii) How do stoichiometric constraints modulate the effects of enrichment and warming on biomass distribution across multiple trophic levels?

## Methods: Population dynamic models

### The Rosenzweig-MacArthur (RM) model

Rates of change of the consumer and resource biomass densities 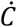 and 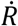 depend on their respective biomass densities *C* and *R* (g.m^−3^):

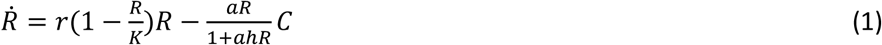

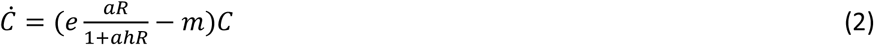

The population growth rate of the resource is given by the logistic equation where *r* is the resource maximum growth rate and *K* is the resource carrying capacity. The population growth rate of the consumer is equal to its feeding rate multiplied by its assimilation efficiency *e* (i.e. the fraction of resource biomass converted into consumer biomass) minus a loss term associated to metabolic losses *m*. The feeding rate of the consumer *C* depends on the density of its resource *R* and follows a Holling type II functional response, with consumer-resource attack rate *a* and handling time *h*.

In the RM model, consumer and resource population growth rates are only limited by nutrient or resource density. Nutrient enrichment is assumed to increase resource carrying capacity, which often leads to the well-known paradox of enrichment where populations fluctuates up to extinctions (Rosenzweig 1971). Nevertheless, this model neither considers nutrient dynamics nor temporal variations of resource stoichiometry and their consequences on population dynamics. To circumvent these limitations of the RM model, we extended it to better consider nutrient dynamics, resource stoichiometry and the way they can affect resource and consumer population dynamics.

### The Stoichiometric Rosenzweig-MacArthur (SRM) model

We derived a stoichiometric extension of the Rosenzweig-MacArthur consumer–resource model with additional stoichiometric and temperature dependencies of several biological rates. We considered two stoichiometric constraints: one on the resource population growth rate, and the other on the consumer assimilation efficiency (see below for more details). These stoichiometric constraints have been observed for several consumer-resource pairs suggesting that they are core components of species growth and interactions (Sterner & Elser 2002).

#### Stoichiometric constraint on the resource population growth rate

Inspired by previous stoichiometric models (Andersen 1997; Loladze *et al.* 2000; Andersen *et al.* 2004; Diehl *et al.* 2005), we extended the RM model by considering explicit nutrient dynamics and nutrient effects on resource population growth rate. The system is assumed to be closed for nutrients. Thus, nutrient supply originates exclusively from biomass excretion and remineralization. The total amount of nutrients in the system (*N*_tot_) is then a measure of nutrient enrichment. In contrast to the very high plasticity in C:N or C:P exhibited by autotrophs, heterotrophs regulate elemental composition within narrower bounds, even when consuming food with large variation in elemental composition (Andersen & Hessen 1991; Sterner & Hessen 1994; Andersen 1997; Elser *et al.* 2000). In other words, the elemental homeostasis is much stronger for consumers compared to primary producers. We thus assumed the nutrient quota (i.e. the nutrient to carbon ratio) of the consumer *Q*_C_ to be conserved whereas the one of the resource *Q*_R_ is flexible over time with the only constraint that Q_R_ > 0. As in the RM model, rates of change of the consumer and resource biomass densities 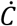 and 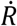 depend on their respective carbon biomass densities *C* and *R* (gC.m^− 3^), except that the resource population growth rate follows the Droop equation (Droop 1974) given by *r*(1−*Q*_min_/*Q*_R_)*R* and is now limited by *Q*_R_ relative to the minimum nutrient quota *Q*_min_ :

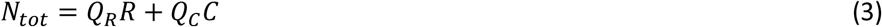

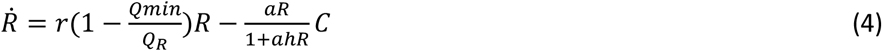

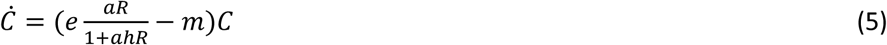

From the nutrient conservation equation (eqn 3) we obtain that 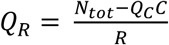. The intuitive interpretation is that the resource nutrient quota *Q*_R_ changes instantaneously with the density of the resource population *R* and with the density of the nutrient stored in the consumer biomass *Q*_C_*C*, to maintain nutrient balance (see Text S1 for details).

### Stoichiometric constraint on the consumer population growth rate

In the RM model, the growth rate of the consumer population only depends on resource density. In other words, the RM model assumes that resource stoichiometry is not limiting and conversion efficiency *e* is often taken for a consumer feeding on a high quality resource (Yodzis & Innes 1992; Binzer *et al.* 2012; Fussmann *et al.* 2014; Uszko *et al.* 2017). However, conversion efficiency can be much lower when the resource is of poor quality (i.e. when there is a stoichiometric unbalance between the consumer and the resource nutrient: carbon ratio) (Elser *et al.* 2000; Elser *et al.* 2007). We relaxed this assumption of the RM model by making the population growth rate of the consumer dependent on both resource quality (i.e. nutrient quota) and quantity (i.e. biomass density). In the SRM model, consumer production is also limited by resource quality as the consumer assimilation efficiency *e* is a saturating function of resource nutrient quota *Q*_R_ :

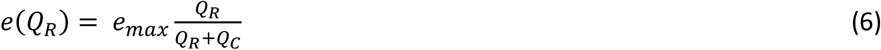

The intuitive interpretation of eqn. 6 is that resource quality is not a limiting factor for consumer growth as long as the nutrient content of the resource is superior to the nutrient content of the consumer (i.e. *Q*_R_ > *Q*_C_). In other words, *e*(*Q*_R_) is proportional to *Q*_R_ for *Q*_R_ < *Q*_C_ and is at its maximum (*e*_max_) for *Q*_R_ > *Q*_C_. The later scenario corresponds to the assumption of the RM model where conversion efficiency is taken for a high quality resource and thus *e* = *e*_max_. By replacing *e* by *e*(*Q*_R_) in eqn. 5, we obtain the SRM model.

### Temperature dependence of model parameters

To investigate the effect of temperature and stoichiometric constraints on consumer-resource dynamics, we next extended the RM and SRM models described above by adding thermal dependencies of the parameters. Following Uszko et al. (2017), we assumed that the total amount of nutrient *N*_tot_, the maximum food conversion efficiency *e*_max_, and fixed stoichiometric traits (*Q*_C_) are independent of temperature, as there is no evidence of systematic temperature dependence for any of them (Peters 1983; Ahlgren 1987; Borer *et al.* 2013; Yvon-Durocher *et al.* 2015). Rate of maintenance respiration and natural background mortality *m* typically increases exponentially with temperature (Fig. S1a and b). We thus used the Arrhenius equation to describe the effect of temperature *T* (in Kelvin) on *m*:

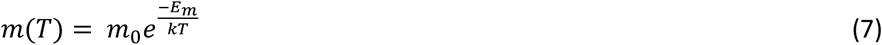

where *m*_0_ is a parameter-specific constant calculated at temperature of 0°C (= 273.15 K). The temperature dependence is characterized by the respective activation energy *E*_m_ (eV) and the Boltzmann constant *k=*8.62×10^−5^ eVK^−1^. As the temperature dependencies of resource intrinsic growth rate *r* and functional response parameters (*a*, 1/*h*) are often unimodal rather than exponential (Englund *et al.* 2011; Rall *et al.* 2012; Sentis *et al.* 2012; Thomas *et al.* 2012), we used Gaussian functions for *r* and *a* and an inverted Gaussian function for *h*:

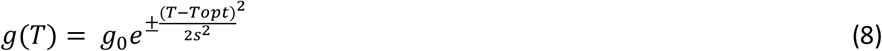

where *T*_opt_ is the temperature at which the rate *g* reaches its minimum or maximum, *s* is the function width and *g*_0_ is a parameter-specific constant calculated at *T*_opt_. The minus-sign corresponds to Gaussian functions and the plus-sign to inverted Gaussian functions.

### Model parameterisation and simulations

To parameterise the models we assumed the resource and consumer species to be a unicellular freshwater algae and a *Daphnia* grazer, respectively. The choice for this system was motivated by the good characterization of both the stoichiometric parameters and thermal dependencies for this system (Andersen 1997; Uszko *et al.* 2017). Uszko *et al.* (2017) recently estimated the thermal dependencies for biological rates of the green algae *Monoraphidium minutum* and the grazer *Daphnia hyalina*. We thus used their estimates of stoichiometric parameters and thermal dependencies (See Table S1 and Fig. S1 for further details).

To investigate the individual and combined effects of enrichment, warming, and stoichiometric constraints, we varied temperature (401 values ranging from 0 to 40°C by 0.1°C) and total amount of nutrients (parameter *N*_tot_ in eqn. 10; 60 values ranging from 0.001 to 0.06 gP.m^−3^ by 0.001 gP.m^−3^, overlapping with reported mean phosphorus concentration in European peri-alpine lakes (Anneville *et al.* 2005)). For the RM model, we used the minimum nutrient quota to convert nutrients into resource (i.e. *K* = *N*_tot_/*Q*_min_). This implies that carrying capacity is independent of temperature which is expected for closed, nutrient-limited systems (Uszko *et al.* 2017) although more experimental evidence are needed to verify this assumption. We then simulated the consumer-resource dynamics for 1000 days to enable the system to reach an attractor (either an equilibrium point or a limit cycle) before we assessed the final state. Therefore, for each model, we simulated 24060 combinations of environmental conditions (401 temperatures by 60 nutrient concentrations). Initial biomass density of each species was set to 0.98 times its equilibrium density in the two-species system (calculated by solving for the two-species equilibrium, using either eqns 1-2 for model RM or eqns 3-5 for model SRM). The value of 0.98 was chosen to be (1) close enough to equilibria to avoid extinctions caused solely by transient dynamics and (2) not exactly the equilibrium value to probe the stability of the equilibrium. Any population falling below the extinction threshold of 10^−9^ g.m^−3^ during the simulation was deemed extinct and its biomass set to zero to exclude ecologically unrealistic low biomass densities. For each model, we calculated system persistence as the percentage of simulations with the two species remaining extant at the end of the simulations. We also calculated system persistence without considering the extinction threshold to assess the proportion of extinctions that are driven by population fluctuations resulting in unrealistic low biomass densities. Population dynamics were simulated with R version 3.4.3 (R Development Core Team 2017) using the “deSolve” package (Soetaert *et al.* 2012) with an absolute error tolerance of 10^−10^ and a relative error tolerance of 10^−6^.

## Results

### Stability: population fluctuations and persistence

Stoichiometric constraints dampened the paradox of enrichment, reducing fluctuations at high nutrient levels and hence increasing persistence. However, stoichiometric constraints also reduced the persistence of the consumer at low and high temperatures. As a result, the overall effect of stoichiometric constraints on stability depends on its relative influence on population fluctuations versus consumer persistence. In the two following paragraphs, we explain in more detail these results and highlight key differences between the outcomes from RM and SRM models.

The RM model predicts that increasing nutrient concentration is strongly destabilizing: the system shifts from a stable equilibrium point to limit cycles (i.e. the system crosses a Hopf bifurcation). This agrees with the paradox of enrichment. As population biomass fluctuations (i.e. cycle amplitude) increase with nutrient concentration, minimal population densities are very low at high nutrient concentrations leading to the extinction of both the consumer and resource once the extinction threshold is crossed (Fig. 1). In the range of temperatures where the consumer persists, warming does not have a strong influence on the nutrient concentration at which the system shifts from the stable equilibrium point to limit cycles, although this qualitative shift is absent at very high temperatures (i.e. 32°C) when the consumer is close to extinction. Warming decreases fluctuation amplitude and thus dampens extinctions driven by the paradox of enrichment, which results in warming enhancing the persistence of the consumer-resource system at high nutrient concentrations. However, very warm and cold temperatures cause the extinction of the consumer (see below for the mechanisms underlying extinctions), releasing resources from top-down control. Overall, we found that, without considering the extinction threshold of 10^−9^ g.m^−3^ (see Model parametrisation and simulations), both the consumer and the resource can persist in 74% of the temperature-nutrient concentration scenarios (i.e. black + orange areas in Fig 1C). Nevertheless, when considering the extinction threshold, they persist in only 21% of the temperature-nutrient scenarios (i.e. black area in Fig. 1c). In other words, comparing the model simulations with and without extinction threshold revealed that, in the RM model, extinctions are mostly driven by population fluctuations leading to very low biomass densities at which the population is at risk of extinction.

**Figure. 1.**
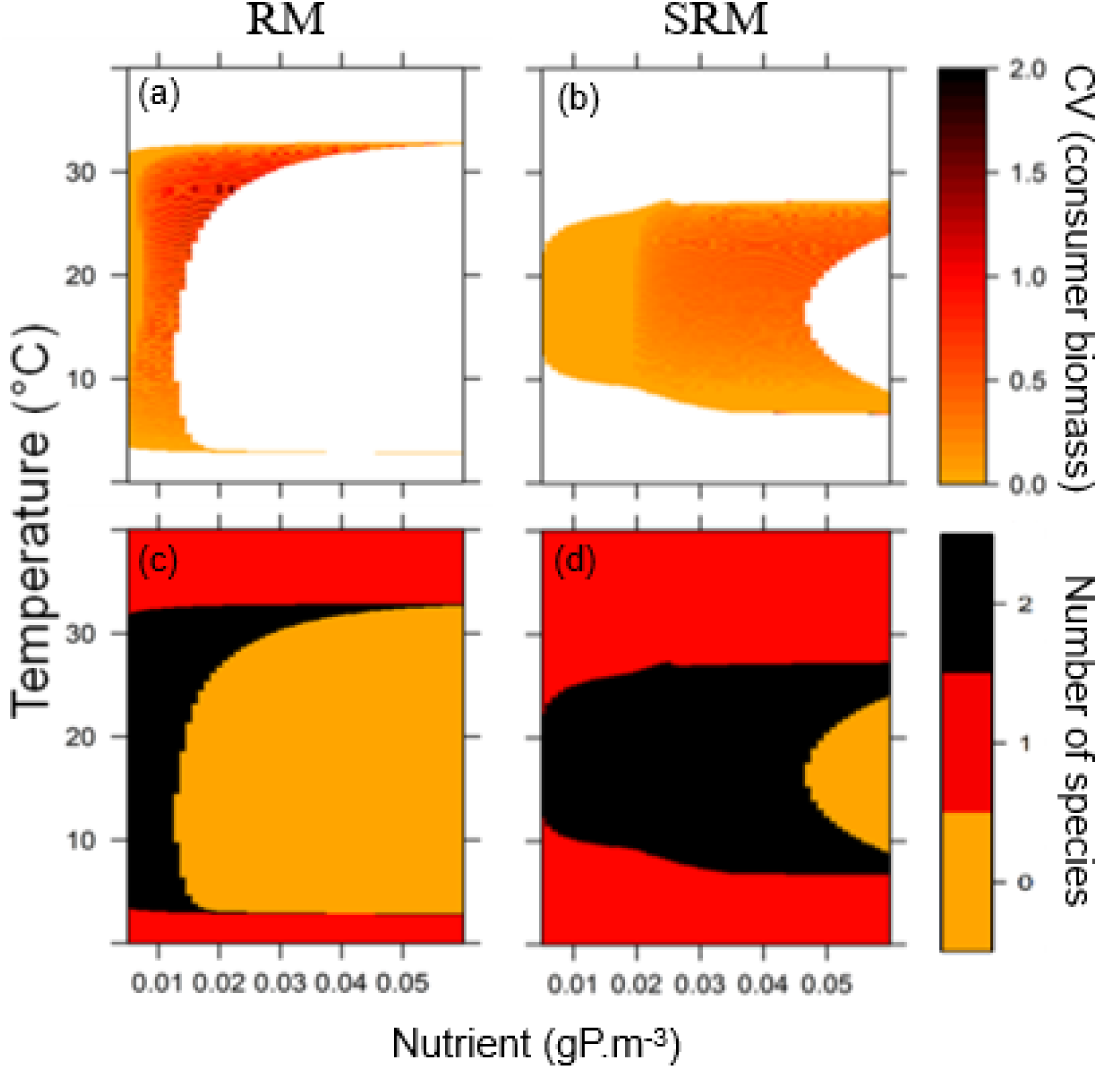
Population fluctuations (consumer biomass coefficient of variation; panels *a* and *b*) and species persistence (number of species; panels *c* and *d*) across the temperature (*y* axis) and nutrient (*x* axis) gradients as predicted by the Rosenzweig-MacArthur (RM; panels *a* and *c*) and by the Stoichiometric Rosenzweig-MacArthur (SRM; panels *b* and *d*) models. In panels *a* and *b*, the white colour corresponds to the temperature-nutrient scenario for which the consumer has gone extinct whereas the orange to red to dark red represent population fluctuations of increasing amplitude. In panels *c* and *d*, in black: both consumer and resource persist; in red: only the resource persists; in orange: none persists. Resource biomass CV is not shown; it is qualitatively similar to the consumer biomass CV as resource and consumer biomass fluctuation are strongly coupled.

In contrast, the SRM model shows that increasing nutrient concentrations causes fewer fluctuations than those observed for the RM model (Fig. 1). This is because: (1) more nutrients are needed to shift the system from a stable equilibrium point to limit cycles—the system can indeed persist without fluctuations up to 0.02 gP.m^−3^ whereas it was only up to 0.0005 gP.m^−3^ in the RM model—and (2) when the system fluctuates, the amplitude of the fluctuations is smaller in the SRM than in the RM model. As a result, stoichiometric constraints dampen the amplitude of population fluctuations (i.e. the paradox of enrichment) and hence increase system persistence at high nutrient levels. While the qualitative effect of temperature is similar to that observed in the RM model, the thermal thresholds for consumer persistence are reduced at low and high temperatures in the SRM predictions. Moreover, thermal thresholds remain almost constant along the nutrient gradient in the RM model, whereas in the SRM model they depend on nutrient concentration, with a smaller thermal range at low nutrient levels compared to high nutrient levels (Fig. 1). The consumer is thus more likely to go extinct at low nutrient concentrations and extreme temperatures in the SRM model than in the RM model. Overall, system persistence for the SRM model was 44% without considering the extinction threshold and 37% when considering it. In other words, comparing the model simulations with and without extinction threshold revealed that, in the SRM model, few extinctions are driven by population fluctuations leading to very low biomass densities. We thus conclude that the RM model predicts larger population fluctuations leading to high probabilities of populations extinctions in comparison to the SRM model.

### Biomass distribution

We next compared the predictions of both models for consumer-resource biomass ratios along the temperature and nutrient gradients (Fig. 2). We found that the RM model systematically predicts biomass ratio > 1 (i.e. consumer biomass is larger than resource biomass). In Contrast, the SRM model predicts biomass ratios both > or < than 1 depending on temperature and nutrient levels. The RM model predicts that, as soon as the consumer can persist, its population biomass density always exceeds the resource population biomass density (Fig. 2). With the SRM model, the biomass ratios are below one at low nutrient levels (Fig. 2). However, at medium and high nutrient levels, the ratios are above one as soon as the consumer can persist. We found qualitatively similar results when considering unstable equilibrium points (Fig. S2). Finally, we showed that, for equivalent parameter values, the RM model predicts biomass ratio that are superior or equal to the ones predicted by the SRM model (text S2). This difference between the two models is independent of the shape and position of the temperature function used to parametrise the models.

**Figure. 2.**
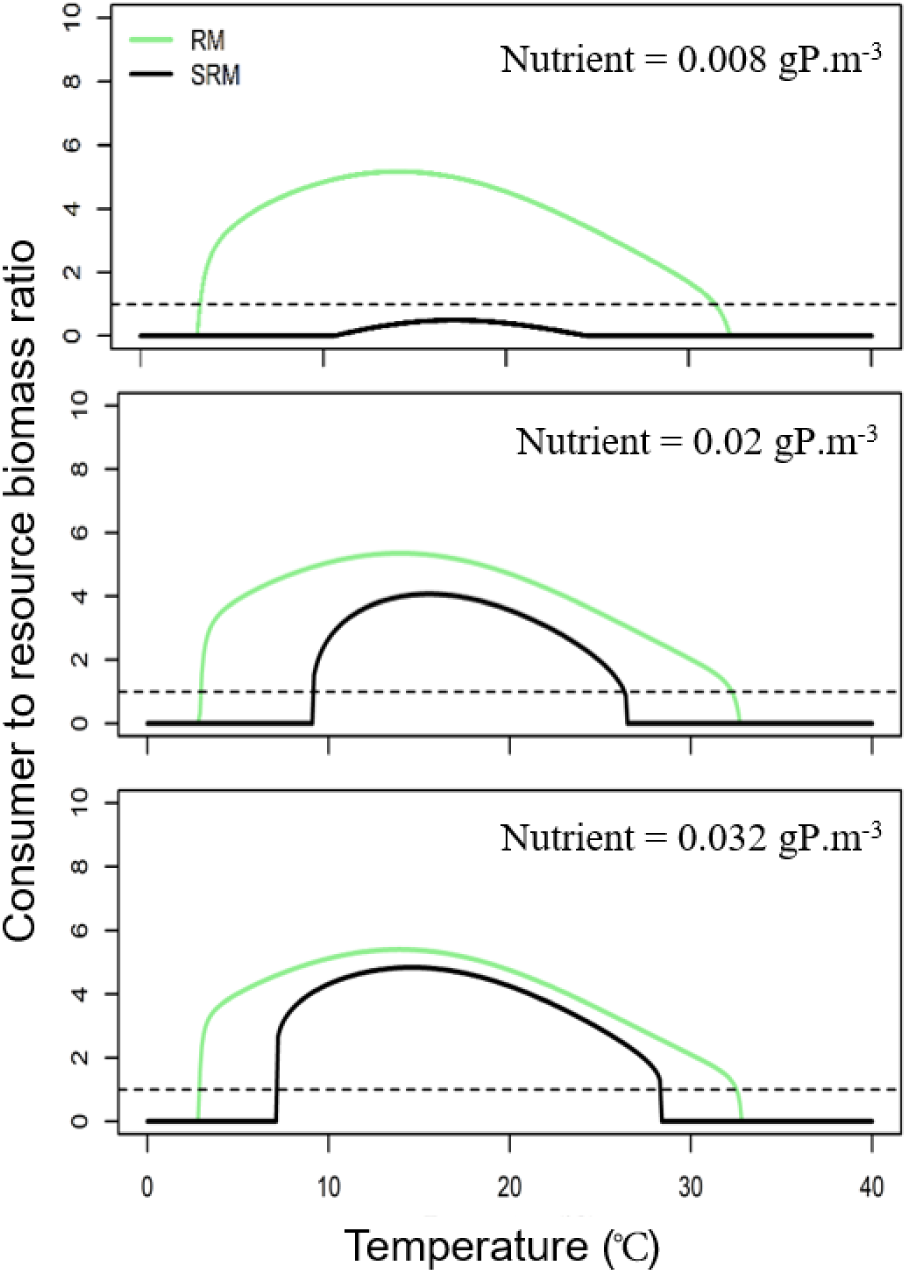
Consumer-resource biomass ratio along the temperature gradient for the Rosenzweig-MacArthur (RM, green lines) and the Stoichiometric Rosenzweig-MacArthur (SRM, black lines) models at three nutrient concentrations (0.008, 0.02, and 0.032 gP.m^−3^). In each panel, the dotted line represents biomass ratio of one; i.e. the biomass densities of the resource and the consumer are equal. Biomass values shown at equilibrium points. For unstable equilibrium points (i.e. limit cycles), see Fig. S2.

### Mechanisms underlying stability and biomass distribution patterns

Here, we detail the mechanisms underlying the stability and biomass distribution patterns to better understand how and when stoichiometric constraints modulate the effects of temperature and nutrients on consumer-resource dynamics. The first mechanism corresponds to the effect of stoichiometric constraints on the consumer energetic efficiency that determines the consumer persistence at extreme low and high temperatures. The second mechanism relates to the influence of the stoichiometric constraints on population dynamical feedback that explains why the stoichiometric model predicts more stability at high nutrient levels compared to the non-stoichiometric model.

### Consumer energetic efficiency

The persistence of the consumer at low and high temperatures is driven by the energetic efficiency *EE* of the consumer (i.e. its feeding rate relative to metabolic losses) calculated as follows:

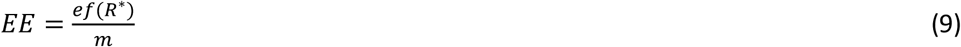

Where *f*(*R*\*) is the functional response of the consumer at resource density *R*\* (i.e. the resource equilibrium density in absence of the consumer). We recall that the assimilation efficiency *e* is a function of resource quality *Q*_R_ in the SRM model whereas it is constant in the RM model. The intuitive interpretation of eqn. 9 is that *EE* should be above one for the consumer population to grow and persist.

To better understand the influence of stoichiometric constraints on consumer persistence, we thus investigated differences in the RM and SRM model predictions regarding the consumer energetic efficiency *EE* along the temperature gradient at two nutrient concentrations (Fig. 3). For both models, energetic efficiency at equilibrium has a hump-shaped relationship with temperature with maximal efficiency values at medium temperatures. While this unimodal shape is conserved across nutrient levels and models, the RM model systematically predicts higher consumer energetic efficiency values than the SRM model because consumer assimilation efficiency is lower in the SRM than in the RM model (Fig. S3). As a result, the temperatures at which energetic efficiency falls below one and drives consumers extinct are more extreme in the RM model compared to the SRM model (Fig. 3). In other words, energetic efficiency is above one for a narrower thermal range in the SRM model.

**Figure. 3.**
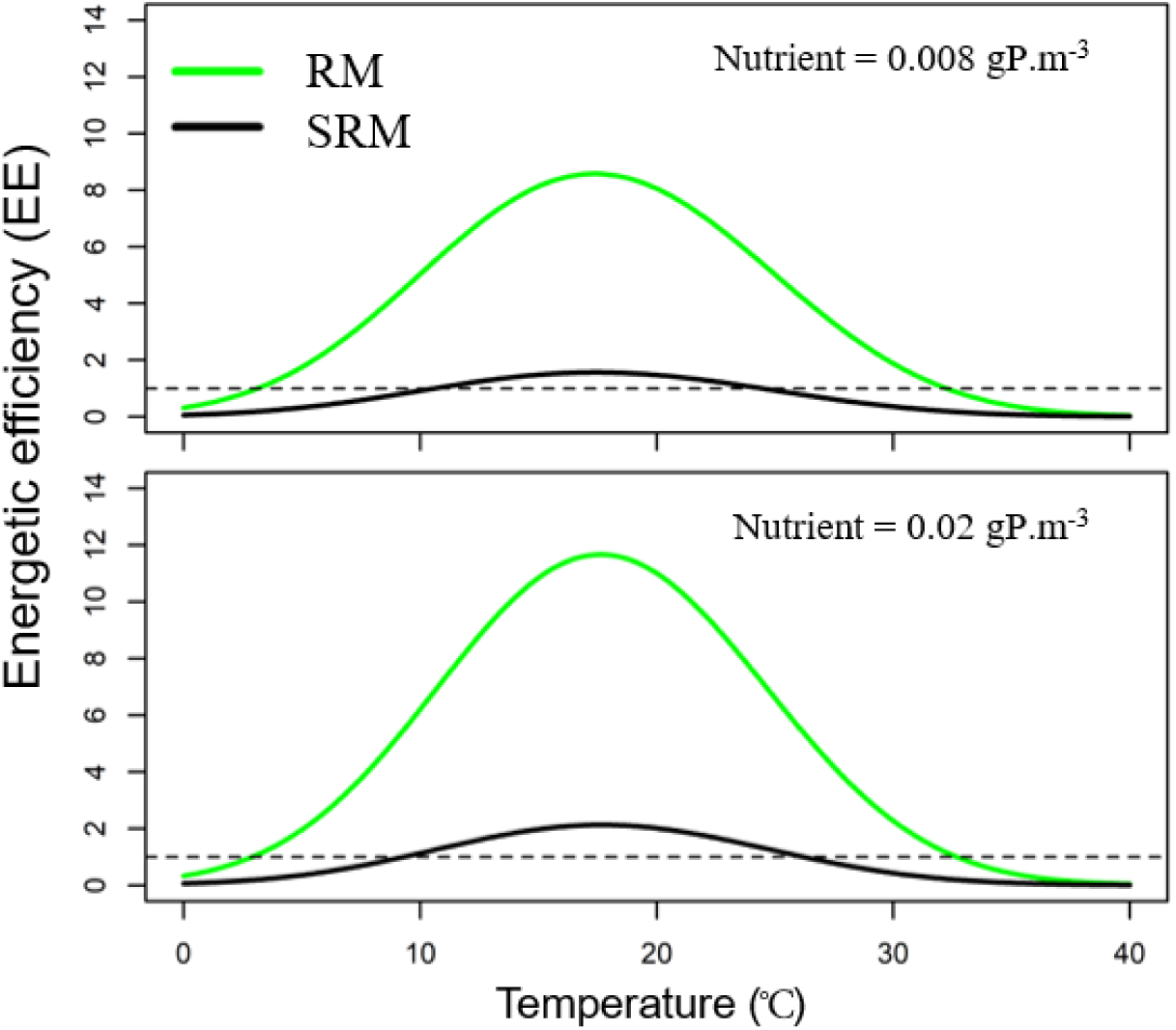
Consumer energetic efficiency along the temperature gradient for the Rosenzweig-MacArthur (RM, in green) and the Stoichiometric Rosenzweig-MacArthur (SRM, in black) models at two nutrient concentrations (0.008 and 0.02 gP/m^3^). In each panel, the dotted line represents energetic efficiency equal to one.

### Dynamical feedbacks due to the stoichiometric constraints

The second mechanism by which stoichiometric constraints influence consumer-resource stability and biomass distribution are the dynamical feedbacks due to stoichiometric constraints on the resource population growth rate and on the consumer energetic efficiency. In the SRM model, the growth rate of the resource population depends on both the total nutrient load and the consumer population density as *Q*_R_ = (*N*_tot_ − *Q*_C_*C*)/*R*. In other words, when consumer population increases, this decreases resource population growth by reducing both resource density (through predation) and quality (through the total nutrient load) leading to a negative feedback on consumer population growth rate. In contrast, for the RM model, the negative consumer feedback is only driven by the reduction in resource density as resource quality is not considered. In addition to this first dynamical feedback, there is a second dynamical feedback as the consumer population growth rate also depends on *Q*_R_ and thus on its own biomass density. Thus, also this second negative feedback loop limits the consumer population growth rate when its density increases. Altogether, dynamical feedbacks reduce strongly the amplitude of population fluctuations, which in turn increases resource and consumer persistence.

To reveal the dynamic effects of the stoichiometric constraints, we calculated the values of assimilation efficiencies and carrying capacities predicted by the SRM model for each temperature-nutrient scenario (Fig. S3) and used these effective parameter values to replace the values of parameters *e* and *K* in the RM model for each temperature-nutrient scenario. In other words, we calculated average values of *e* and *K* in the dynamic SRM model and used them as constant input parameters in the RM model. The objective of using these effective parameter values was to disentangle the static effect of stoichiometric constraints (i.e. changing the average parameter values of consumer assimilation efficiency and of the resource carrying capacity) from their population dynamical effect (i.e. the two dynamical feedback described above). We thus simulated population dynamics along the temperature-nutrient gradient using the RM model with these effective parameters; referred hereafter as effective RM model (Fig. 4). Comparing predictions from the RM, effective RM, and SRM models allowed to disentangle the static stoichiometric effects when going from the RM to the effective RM predictions (Fig. 4, panels a to b) from the dynamical stoichiometric effects when going from the effective RM to the SRM predictions (Fig. 4, panels b to c). In other words, the RM and effective RM only differ in their parameter values because the effective RM takes into account the effect of stoichiometric constraints on the average parameter values. On the other hand, the effective RM and SRM have similar parameter values but different population dynamics, which helps understanding the dynamical feedback induced by stoichiometric constraints.

**Figure. 4.**
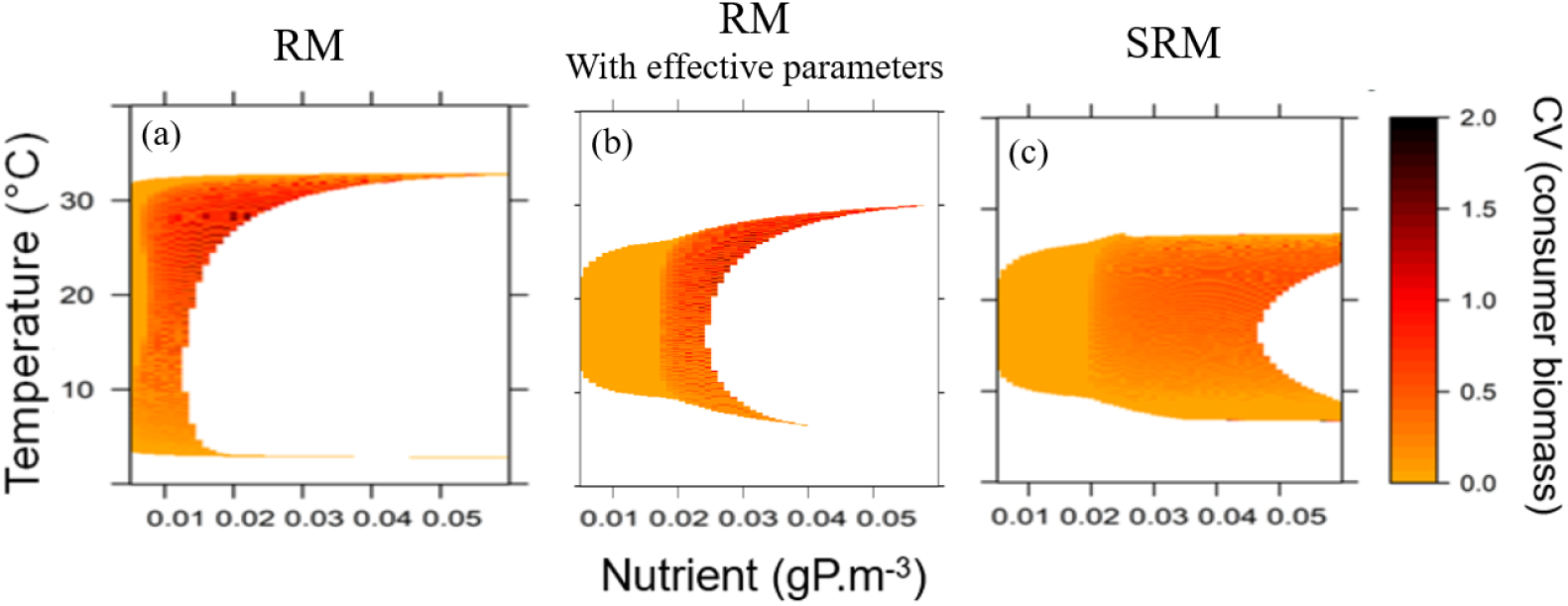
Population fluctuations (consumer biomass coefficient of variation) across the temperature (*y* axis) and nutrient (*x* axis) gradients as predicted by the Rosenzweig-MacArthur (RM; panel *a*), the RM with effective parameters (panel *b*), and the Stoichiometric Rosenzweig-MacArthur (SRM; panel *c*) models.

We found that, at low nutrient concentrations, population fluctuations and consumer persistence predicted by the effective RM model agreed with predictions of the SRM model. However, the system shifted from a stable equilibrium point to a limit cycle at lower nutrient concentrations for the effective RM model than for the SRM model. This suggests that more nutrients are needed to destabilize the system with the SRM model. Moreover, the effective RM model predicts ampler population fluctuations than the SRM model. As a result, the effective RM predicts high extinction rates at high nutrient concentrations compared to the SRM model. Overall, we found that the effective RM model cannot fully reproduce the dynamics predicted by the SRM, which indicates that including stoichiometric constraints in the RM model involves more than only changing parameter values.

## Discussion

Temperature and nutrient enrichment are two of the most important drivers of global change (Nelson 2005). However, most research on the effects of temperature and nutrients on community dynamics assumes that the elemental composition of primary producers and consumers are constant and independent of changes on energy and material fluxes (Binzer *et al.* 2012; Boit *et al.* 2012; Amarasekare & Coutinho 2014; Gilbert *et al.* 2014; Amarasekare 2015; Binzer *et al.* 2016; Gilarranz *et al.* 2016). Yet, the elemental composition of primary producers is known to be flexible, which can have important consequences for community dynamics and ecosystem processes (Elser *et al.* 2000). We have shown how stoichiometric constraints that account for flexible stoichiometry can affect predictions on how temperature and nutrients influence community stability and biomass distribution across trophic levels. We thus argue that considering stoichiometric constraints is an important step toward a better understanding of the effects of global change on ecosystems.

### Stoichiometric constraints and temperature can dampen the paradox of enrichment

We showed that both stoichiometric constraints and temperature dampen the negative effect of nutrient enrichment on consumer-resource fluctuations and increase system persistence at high nutrient levels. Temperature effects are driven by physiological mechanisms. In agreement with previous empirical studies, our model parametrization reflects the observation that metabolic loss rates increase faster with warming than consumer feeding rates (Vucic-Pestic *et al.* 2011; Sentis *et al.* 2012; Fussmann *et al.* 2014; Iles 2014). Consumers are thereby less energetically efficient at higher temperatures which stabilizes food-web dynamics by reducing energy flow between trophic levels (Binzer *et al.* 2012; Kratina *et al.* 2012; Fussmann *et al.* 2014; Sentis *et al.* 2017). In contrast, the effect of stoichiometric constraints is mainly linked to two mechanisms: a shift in the position of the Hopf bifurcation and negative dynamical feedbacks of the consumer and resource on their population growth rates. Both resources and consumers are composed of the same essential elements (N, P, and C), which implies that, when consumer or resource population biomass increases, it reduces the pool of free nutrients available for the growth of the resource population. Therefore, more nutrients are needed to shift the system from a stable equilibrium to population cycles. In other words, the paradox of enrichment is displaced to higher nutrient concentrations (i.e., the position of the Hopf bifurcation is shifted to higher nutrient levels. In contrast, the RM model does not take into account the storage of nutrients in both the resource and consumer biomasses (i.e. the carrying capacity only depends on the total nutrient load). Less enrichment is thus required to shift the system from a stable equilibrium point to limit cycles.

We found two dynamic effects that correspond to negative dynamical feedbacks of the consumer and the resource on themselves. When consumer population increases, it decreases the population growth rate of the resource by limiting nutrient availability, diminishing resource biomass, which, in turn, decreases the consumer population growth rate. Conversely, when the resource biomass increases, this decreases the nutrient content of the resource, which, in turn, limits the growth rates of both the resource and consumer populations. These stoichiometric negative feedback loops strongly decrease the amplitude of population fluctuations and thus dampen the paradox of enrichment. Interestingly, our comparisons of the RM, effective RM and SRM model predictions indicate that the dynamical effects contribute more to the reduction of fluctuations than the static effects: population fluctuations are large in the effective RM model accounting for the static effect only, whereas they are much smaller in SRM model accounting for both static and dynamical effects (Fig. 4). This implies that the impact of stoichiometric constraints on community dynamics goes beyond a simple modification of parameter values and encompass more complex population feedbacks between the consumer and the resource.

Overall, these results demonstrate that both flexible stoichiometry and temperature can synergistically dampen the paradox of enrichment by two different mechanisms: population dynamic feedbacks and physiological constraints. Our consumer-resource model is simplified compared to natural communities composed of numerous species. Moreover, in natural systems, a large amount of nutrient can be stored in abiotic and slow biotic pools that have long turnover times which, in turn, can influence the population dynamics. In particular, the amplitude of the population fluctuations is expected to be smaller as abiotic pools can buffer the population feedback. Nevertheless, considering the nutrient held in slow abiotic or biotic pools would not change the equilibrium densities of primary producers and grazer if nutrients are released in the environment proportionally to their density stored in the abiotic pool (Menge *et al.* 2012). Moreover, the predictions of the stoichiometric model fit with empirical observations. In eutrophic lakes and experimental mesocosms, populations can persist at relatively high nutrient concentrations even if fertilisation enhance population fluctuations (O’Connor *et al.* 2009; Boit *et al.* 2012; Kratina *et al.* 2012), as our stoichiometric model predicts. In contrast, the Rosenzweig-MacArthur model tends to produce very large population fluctuations and extinctions at low nutrient concentrations which can explain why these predictions are not well supported by empirical observations (McAllister *et al.* 1972; Jensen & Ginzburg 2005).

### Effects of stoichiometric constraints on system persistence across environmental gradients

While stoichiometric constraints dampen the paradox of enrichment and thus increase persistence at high nutrient levels, they also reduce the persistence of the consumer at low and high temperatures. Stoichiometric constraints affect the thermal thresholds for consumer extinctions. Consumers can only persist over a narrower range of intermediate temperatures when they are constrained by stoichiometry. This is due to the reduced biomass assimilation of the consumer at low and high temperatures that, in turn, decreases its energetic efficiency and thus fastens consumer extinction. In our stoichiometric model, the reduction of biomass assimilation efficiency emerges from the effect of temperature on resource quality: extreme high and low temperatures decrease resource quality and thus less resource biomass can be converted in consumer biomass at these temperatures. The emergence of a thermal dependency for assimilation efficiency contrasts with previous theoretical studies that used the RM model and assumed that the assimilation efficiency is temperature independent as resource quality is assumed constant (Binzer *et al.* 2012; Gilbert *et al.* 2014; Sentis *et al.* 2017; Uszko *et al.* 2017). In the SRM model, the thermal dependency of the consumer assimilation efficiency is fully driven by the change in the resource stoichiometry induced by temperature. The SRM model thus predicts an additional mechanism by which temperature can influence trophic interactions: temperature changes resource stoichiometry, which, in turn, impacts the consumer assimilation efficiency and its population growth rate. This prediction matches with empirical results showing that primary producer stoichiometric composition can change with temperature (Woods *et al.* 2003) and that consumer assimilation efficiency is sensitive to resource stoichiometric composition (Andersen 1997; Elser *et al.* 2000). To sum up, the overall effect of stoichiometric constraints on system persistence thus depends on the temperature range considered and on their relative influence on population fluctuations versus consumer persistence.

### Effects of stoichiometric constraints on biomass distribution

We found that stoichiometric constraints can modulate the effects of temperature and nutrients on biomass distribution across trophic levels. Without stoichiometric constraints (i.e. with the Rosenzweig-MacArthur model), biomass ratios are above one for almost all temperatures or nutrient levels as the biomass produced by the resource is efficiently transferred to the consumer level consistently along the environmental gradients. This finding agrees with theoretical studies reporting that Lotka-Volterra and RM models predict biomass ratios above one and fail to reproduce biomass pyramids for a substantial region of parameter values (Jonsson 2017; Barbier & Loreau 2019). However, in nature, consumer-resource biomass ratios are often below one (McCauley & Kalff 1981; Del Giorgio & Gasol 1995; McCauley *et al.* 1999; Irigoien *et al.* 2004) suggesting that additional mechanisms should be included to better understand and predict biomass distribution patterns in natural food webs. Our stoichiometric model agrees with experimental observations. It predicts that, at low nutrient concentrations (i.e. < 0.01 gP.m^−3^), the biomass ratio never exceeds one along the entire temperature gradient. This is observed in oligotrophic aquatic systems where primary production is too low to sustain high consumer populations (O’Connor *et al.* 2009). In addition, we also found that increasing nutrient levels decreased the temperature ranges within which biomass ratio is below one. This corresponds to results from manipulated nutrient concentrations and temperature in aquatic mesocosms, where zooplankton to phytoplankton biomass ratio only exceeds one in the enriched mesocosms at medium or warm temperatures (i.e. 27°C) (O’Connor *et al.* 2009). This suggests that the models with stoichiometric constraints better reproduce the biomass patterns observed in experimental and natural systems. Nevertheless, further experiments investigating the links between stoichiometric flexibility and consumer-resource dynamics are needed to determine if these stoichiometric mechanisms are underlying patterns of biomass distribution in nature.

### Implications of our findings for global change

Temperature and nutrients do not act in isolation from each other. Climate warming, for example, causes stronger water stratification which, in turn, can limit nutrient cycling (Sarmiento *et al.* 2004; Tranvik *et al.* 2009). Environmental policies such as the European water framework directive (i.e. Directive 2000/60/EC of the European Parliament and of the Council establishing a framework for the Community action in the field of water policy) effectively reduces input of nutrients in aquatic ecosystems (Anneville *et al.* 2005) while the climate keeps warming. With these two phenomena, water will often be warmer and contain fewer nutrients in aquatic systems. Our models consistently predict that warmer temperatures should stabilise consumer-resource dynamics but, if temperature further increases, the consumer goes extinct as energetic efficiency decreases with warming. Moreover, we found that stoichiometric constraints can reduce this thermal extinction threshold (i.e. the consumer persists in a narrower thermal range), especially at low nutrient levels. Our stoichiometric model thus suggests that decreasing nutrient concentrations alongside warmer temperatures should fasten the extinction of consumer populations. This prediction matches empirical observations of consumer extinctions at warm temperatures in oligotrophic aquatic systems (Petchey *et al.* 1999; O’Connor *et al.* 2009). Altogether, these results indicate that considering stoichiometric constraints can be of importance for the management of nutrient inputs and the conservation of natural populations and communities under climate change.

## Conclusion

Knowledge of how temperature and nutrient simultaneously influence the elemental composition of primary producers and consumers is crucial to better understand and predict the effects of global change on species interactions, community dynamics and fluxes of energy and material within and among ecosystems. Here we showed that stoichiometric constraints dampen the negative effect of enrichment on stability by reducing population fluctuations through population dynamics feedbacks. However, stoichiometric constraints also decrease consumer energetic efficiency, which increases consumer extinction risk at extreme temperatures and low nutrient concentrations. Finally, stoichiometric constraints can reverse biomass distribution across trophic level by modulating consumer efficiency and resource population growth rate along the temperature and nutrient gradients. Our study provides a first step in the exploration of the consequences of stoichiometric constraints and temperature on ecological communities. It suggests that accounting for stoichiometric constraints can strongly influence our understanding of how global change drivers impact important features of ecological communities such as stability and biomass distribution patterns.

## Data accessibility

R codes are available on Dryad (https://doi.org/10.5061/dryad.msbcc2fv2).

## Acknowledgements

We thank Elisa Thébault and two anonymous reviewer for their constructive comments on the manuscript. This work is funded by the FRAGCLIM Consolidator Grant (number 726176) to Jose M. Montoya from the European Research Council under the European Union’s Horizon 2020 Research and Innovation Program. Version 7 of this preprint has been peer-reviewed and recommended by Peer Community In Ecology (https://doi.org/10.24072/pci.ecology.100039).

## Author contributions

A.S., B.H., and J.M.M. conceived the study. B.H. and A.S. developed and analysed the models. A.S. wrote the first draft of the manuscript. All authors contributed substantially to revisions.

## Appendix

### Supplementary information

#### Text S1. Derivation of the Stoichiometric Rosenzweig-MacArthur (SRM) model

The model studied in the main text is very similar to previous stoichiometric consumer-resource models (Andersen 1997; Loladze *et al.* 2000; Andersen *et al.* 2004; Diehl *et al.* 2005). To make our paper self-contained, we here present the model assumptions and derive the model equations (eqs. 3-6 in main text). Our objective was not to develop a complex and very realistic stoichiometric model that would include additional important abiotic and biotic features such as light intensity (Diehl 2007) or compensatory feeding (Cruz-Rivera & Hay 2000). Instead, we aimed at introducing two fundamental stoichiometric features (i.e. stoichiometric flexibility and stoichiometric imbalance) and investigate how these stoichiometric considerations can change predictions of the Rosenzweig-MacArthur model. We assumed that resource and consumer production are limited by energy and a single mineral nutrient. Moreover, we assume the system is closed for nutrients. Thus, nutrient supply originates exclusively from excretion and remineralization of biomass. The total amount of nutrients in the system (*N*_tot_) is then a measure of nutrient enrichment. As elemental homeostasis is much stronger for consumers compared to primary producers (Andersen 1997), we assumed the nutrient quota of the consumer *Q*_C_ to be constant whereas the nutrient quota of the resource *Q*_R_ is flexible. Four differential equations determine the dynamics of four state variables, that is, the concentrations of resource (*R*) and consumer (*C*) carbon biomasses and of dissolved mineral nutrients (*N*), and the nutrient quota of the resource (*Q*_R_):

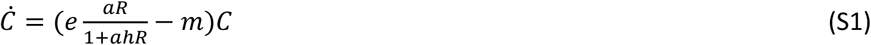

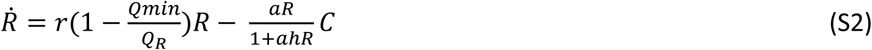

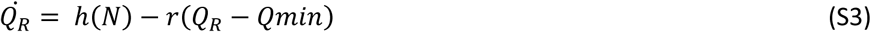

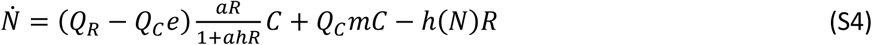

As in the RM model, rates of change of the consumer and resource biomass densities 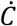 and 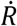 depend on their respective carbon biomass densities *C* and *R* (gC.m^−3^), except that the resource population growth rate follows the Droop equation (Droop 1974) and is now limited by its nutrient quota *Q*_R_ relative to the minimum nutrient quota *Q*_min_. Rate of change of *Q*_R_ depends on the nutrient uptake rate by the resource species *h*(*N*) and the amount of nutrient invested in growth (eqn S3). *h*(*N*) is the specific resource nutrient uptake rate and can be represented by a Michaelis-Menten model where the amount of nutrient uptake saturates at high nutrient concentrations.

With the mass-balance equation, we get that the total amount of nutrient is the sum of the free nutrient plus the nutrient fixed in the resource biomass plus the nutrient fixed in the consumer biomass: *N*_tot_ = *N* + *Q*_R_ *R* + *Q*_C_ *C.* As Eqns S1-S4 conserve total biomass (the system is closed), the time derivative of *N*_tot_ is zero. We can thus replace one of the four differential equations S1-S4 with the algebraic equation *N*_tot_ = *N* + *Q*_R_ *R* + *Q*_C_ *C:*

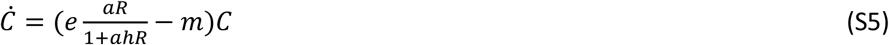

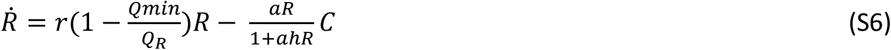

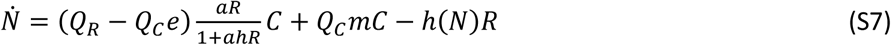

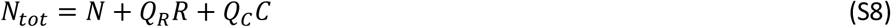

It is possible to derive a simpler model by reducing the number of dimensions in the above model from three to two. This model reduction is based on the assumption that free nutrients are taken up very quickly relative to the dynamics of the consumer and resource biomasses. This corresponds to taking *h*(*N*) large, 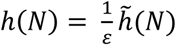 for small ε.

The fast dynamics (on the timescale t ∼ ε) are

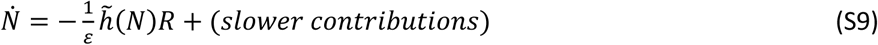

Which converge to N → 0, and 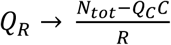 with *N*_tot_ the total nutrient in the system. In other words, *N* in dead and excreted matter is immediately recycled and acquired by the resource species. When substituting the quasi-steady-state in eqns. (S5, S6), we get the resulting dynamics (on the timescale t ∼1):

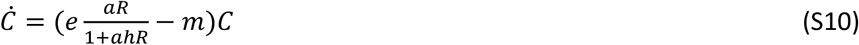

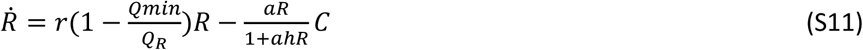

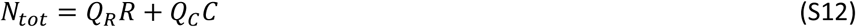

From the nutrient conservation equation (eqn. S12) we obtain that 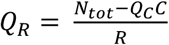. The intuitive interpretation is that the resource nutrient quota *Q*_R_ decreases with the density of the resource population and with the density of nutrient stored in the consumer biomass. In contrast to eqns S5-S8, the reduced model has only two differential equations and one algebraic equation. It can be equivalently written as a set of three differential equations with 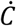 and 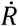 similar as equations S10 and S11 and with 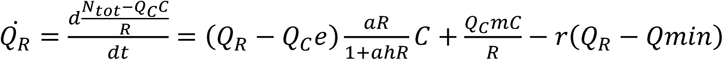.

In the RM model, the growth rate of the consumer population is assumed to depend only on resource density. We relaxed this assumption by making the population growth rate of the consumer dependent on both the resource quality (i.e. nutrient quota) and quantity (i.e. density). In the SRM model, consumer production is also limited by resource quality as the consumer assimilation efficiency *e* is a saturating function of resource nutrient quota *Q*_R_ :

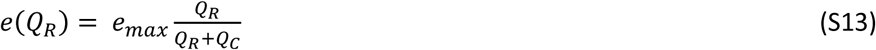

The intuitive interpretation of eqn. S13 is that resource quality is not a limiting factor for consumer growth as long as the nutrient content of the resource is superior to the nutrient content of the consumer (i.e. *Q*_R_ > *Q*_C_). In other words, when *Q*_R_ >> *Q*_C_, *e*(*Q*_R_) → *e*_max_ and when *Q*_R_ << *Q*_C_, *e*(*Q*_R_) → 0. By replacing *e* by *e*(*Q*_R_) in eqn. S10, we obtain the SRM model.

#### Text S2. Differences in biomass ratios predicted by the two models

Here we show that the equilibrium consumer-to-resource biomass ratio in the model with stoichiometric constraints (SRM model) is always smaller than the one in the model without stoichiometric constraints (RM model), keeping the same parameter values. For simplicity we assume for both models that the consumer and the resource persist at equilibrium, and we do not consider the stability of the equilibrium point (in particular, the equilibrium might be unstable at the center of a limit cycle). We indicate the equilibrium values of the non-stoichiometric model by the superscript “ns” and the equilibrium values of the stoichiometric model by the superscript “s”. We use the same superscripts to distinguish the assimilation efficiencies of both models.

##### Model without stoichiometric constraints

The model is defined as

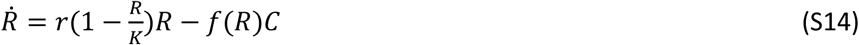

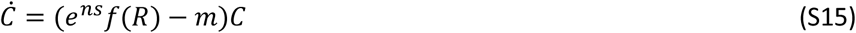

With 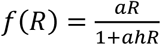 and *K* = *N*_tot_/*Q*_min_.

By solving equation (S15) we get the resource biomass at equilibrium:

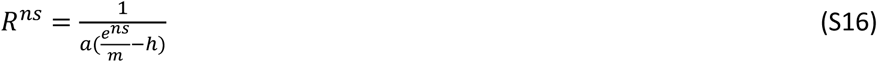

From equation (S14) we get the consumer biomass at equilibrium. It follows from

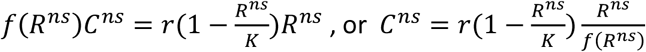

Hence, the consumer-to-resource biomass ratio is

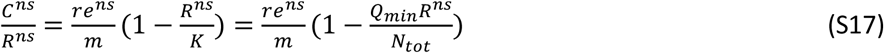

##### Model with stoichiometric constraints

The model is defined as

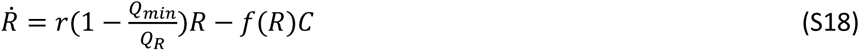

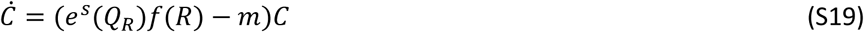

With 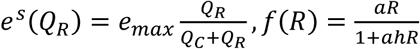 and *N* _*tot*_ *= Q*_*R*_*R + Q*_*C*_*C.*

From equation (S19) we have 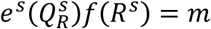, or

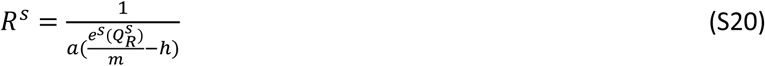

From equation (S18) we have from 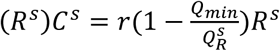, or

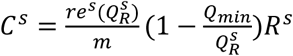

Hence, the consumer-to-resource biomass ratio is

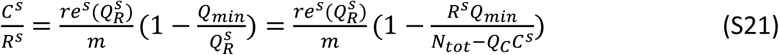

We now compare the biomass ratios of equations (S17) and (S21). We have *e*^ns^ = *e*_max_ as the RM model assumes that resource stoichiometry is not limiting and conversion efficiency is thus at its maximal value. However, conversion efficiency can be much lower when the resource is of poor quality (i.e. when there is a stoichiometric unbalance between the consumer and the resource nutrient: carbon ratio) (Elser *et al.* 2000; Elser *et al.* 2007). In other words, the consequence of stoichiometric constraints is to lower the values of conversion efficiency from the RM model. We thus obtain:

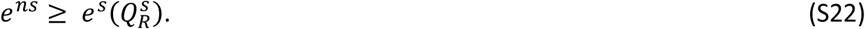

Using this inequality, we get 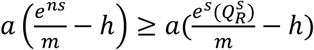, and by equations (S16) and (S20), we see that

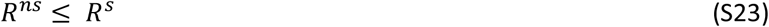

Clearly, we always have *N*_*tot*_ ≥ *N*_*tot*_ − *Q*_*C*_ *C*^*s*^. Combining this with equation (S23), we get 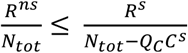 and

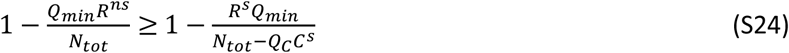

Finally, from equations (S22) and (S24),

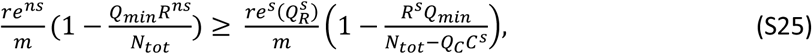

showing that 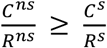.

**Table S1.** Definitions and units of model parameters, from Uszko *et al.* (2017). For temperature-dependent parameters, we list the value of the scaling constant *Q*_0_ (in units of the parameter) and the values of either the activation energy *E*_Q_ (eV, when temperature dependence is monotonous, eqn. 7) or of the temperature *T*_opt_ (Kelvin) at which the parameter value reaches a maximum/minimum and the width *s* (Kelvin) of this bell-/U-shaped function (when temperature dependence is non-monotonous, eqn. 8). Biomass and nutrients are expressed in units of carbon (C) and phosphorus (P), respectively

| Parameter | Temperature independent value | Unit | Definition | Reference |
| --- | --- | --- | --- | --- |
| **Thermal parameters** |  |  |  |  |
| $r$ | $r_0 = 2.2$; $T_{opt} = 298.15$; $s = 12.0$ | 1/d | Intrinsic rate of resource net production (gross production – biosynthesis costs) | Uszko et al. 2017 |
| $h$ | $h_0 = 0.17$; $T_{opt} = 294.1$; $s = 6.4$ | d | Handling time | Uszko et al. 2017 |
| $a$ | $a_0 = 8.9$; $T_{opt} = 296.0$; $s = 9.4$ | $m^3/(gC\ d)$ | Attack rate | Uszko et al. 2017 |
| $m$ | $m_0 = 4.4 \times 10^8$; $E_m = 0.55$ | 1/d | Consumer mortality plus maintenance rate | |
| $e_{max}$ | 0.385 | - | Maximum assimilation efficiency | Peters 1983 |
| $Q_C$ | 0.042 | g P/g C | Consumer P:C ratio | Diehl 2005 |
| $Q_{min}$ | 0.009 | g P/g C | Minimum nutrient quota | Diehl 2005 |
| $Q_R$ | Variable | g P/g C | Resource P:C ratio | |
| $N_{tot}$ | Variable | g P/m<sup>3</sup> | Total nutrients in the system | |
| $T$ | Variable | K | Temperature | |

**Fig. S1.**
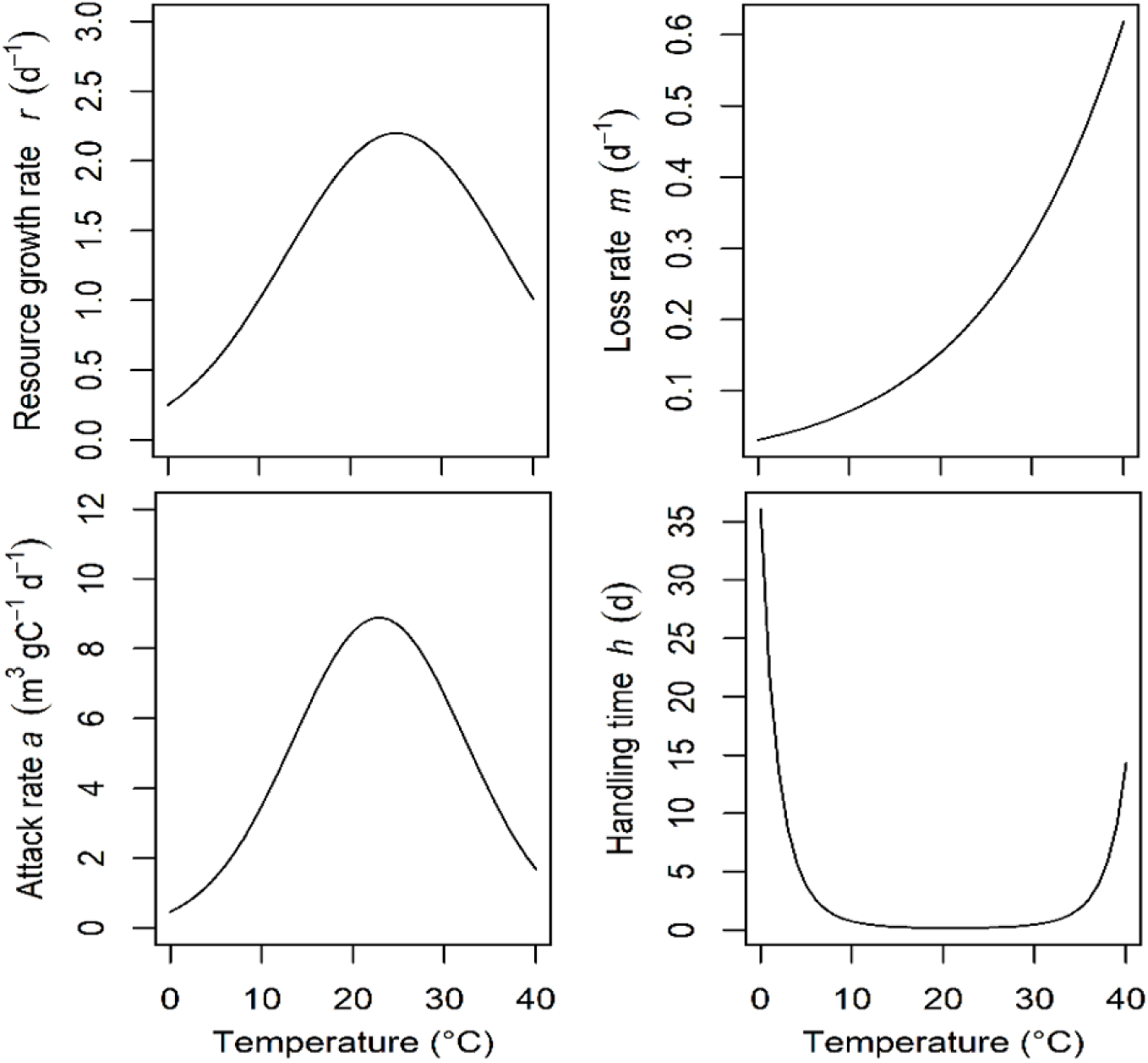
Thermal functions used to parametrize the model (adapted from Uszko et al. 2017)

**Fig. S2.**
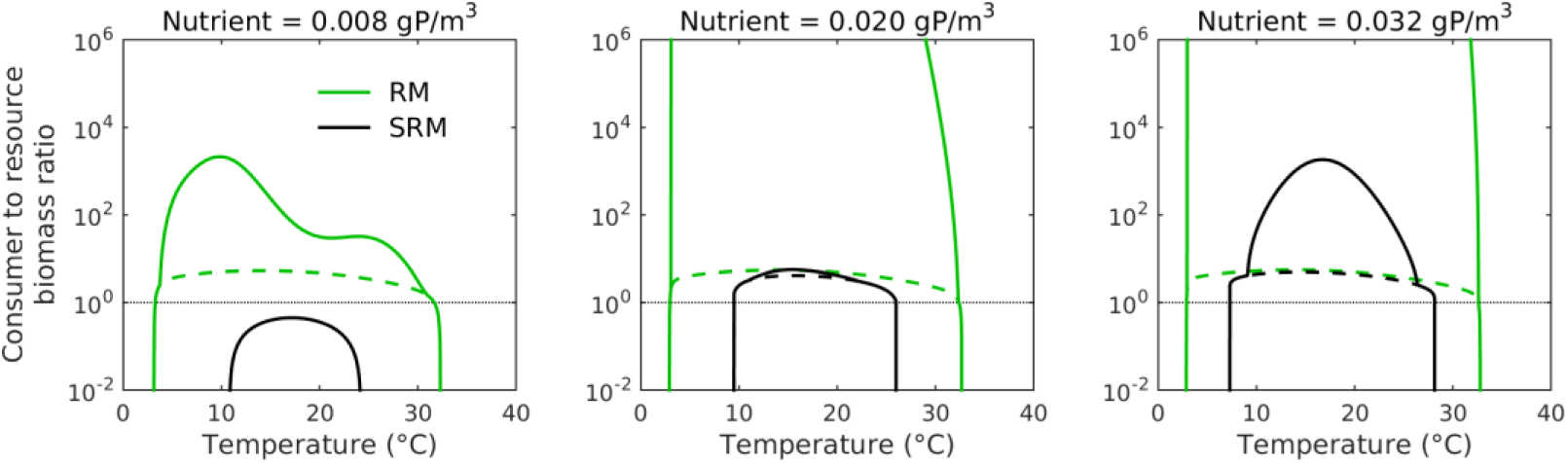
Consumer-resource biomass ratio (log scale) along the temperature gradient for the Rosenzweig-MacArthur (RM, green lines) and the Stoichiometric Rosenzweig-MacArthur (SRM, black lines) models at three nutrient concentrations (0.008, 0.02, and 0.032 gP.m^−3^). In each panel, the dotted lines represent unstable solutions whereas full lines represent stable solutions. The thin horizontal dotted line represents biomass ratio of one; i.e. the biomass densities of the resource and the consumer are equal.

**Fig. S3.**
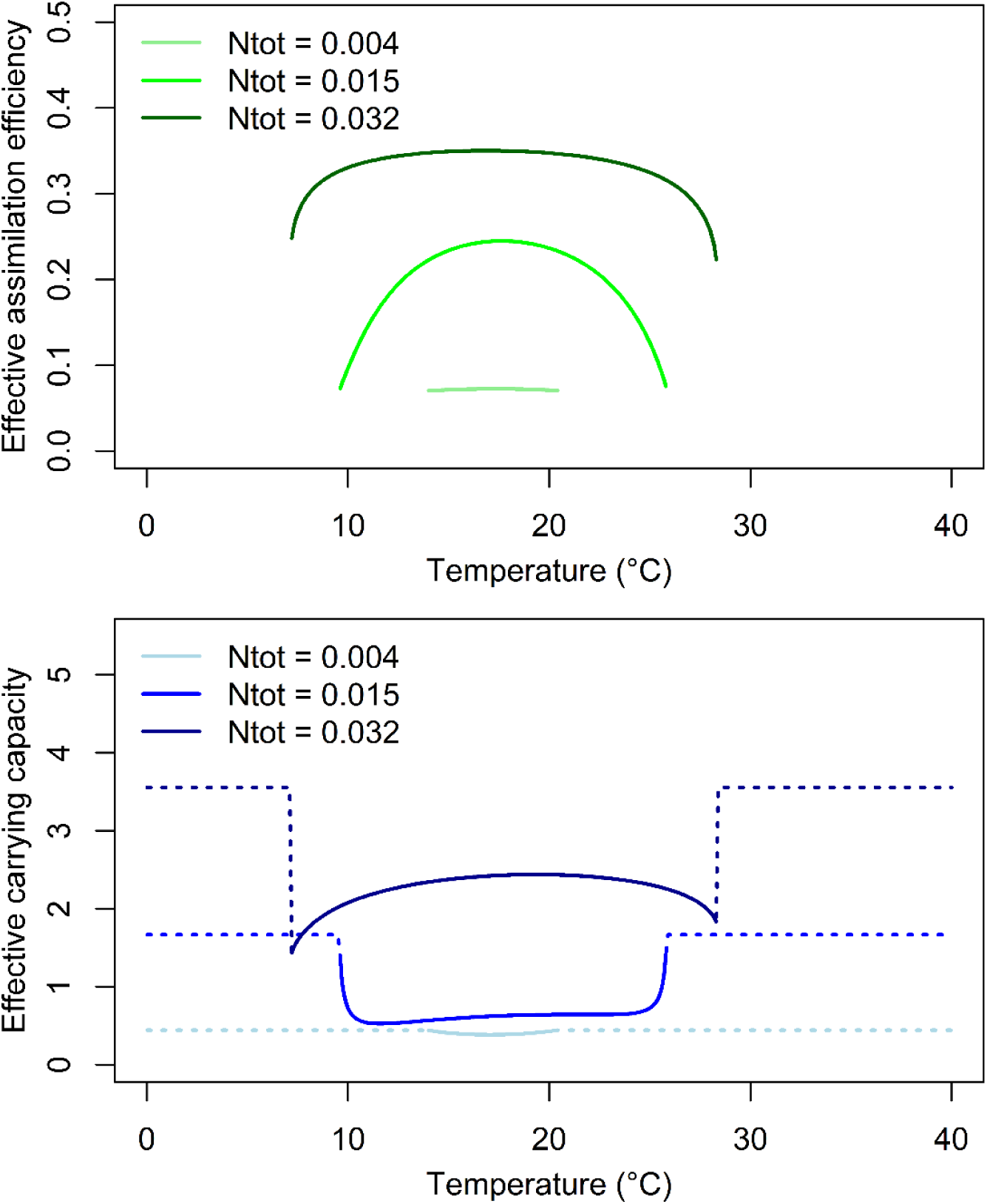
Effective assimilation efficiency *e*_ef_ and carrying capacity *K*_ef_ from the Stoichiometric Rosenzweig-MacArthur (SRM) model along the temperature gradient at three nutrient levels (0.004, 0.015, and 0.032 gP.m^−3^) with *Q*_C_ = 0.042. Full lines represent temperature and nutrient scenarios for which both the resource and consumer persist whereas dotted lines represent scenarios for which only the resource persists. Effective assimilation efficiency was calculated as *e*_ef_ = *e*_max_ *Q*_R_/(*Q*_R_ +*Q*_C_), with *Q*_R_ the equilibrium solution of the SRM model and the effective carrying capacity as *K*_ef_ = *Q*_R_ *R*/*Q*_min_ = (*N*_tot_ − *Q*_C_ *C*)/*Q*_*min*_, with *Q*_R_, *R* and *C* the equilibrium solutions of the SRM model.

## References

Ahlgren, G. (1987). Temperature functions in biology and their application to algal growth constants. Oikos, 49, 177–190.

Amarasekare, P. (2015). Effects of temperature on consumer-resource interactions. Journal of Animal Ecology, 84, 665–679.

Amarasekare, P. & Coutinho, R.M. (2014). Effects of temperature on intraspecific competition in ectotherms. The American Naturalist, 184, E50–E65.

Andersen, T. (1997). Pelagic nutrient cycles: herbivores as sources and sinks. Springer-Verlag, Berlin, Germany.

Andersen, T., Elser, J.J. & Hessen, D.O. (2004). Stoichiometry and population dynamics. Ecology Letters, 7, 884–900.

Andersen, T. & Hessen, D.O. (1991). Carbon, nitrogen, and phosphorus content of freshwater zooplankton. Limnology and Oceanography, 36, 807–814.

Anneville, O., Gammeter, S. & Straile, D. (2005). Phosphorus decrease and climate variability: mediators of synchrony in phytoplankton changes among European peri-alpine lakes. Freshwater Biology, 50, 1731–1746.

Barbier, M. & Loreau, M. (2019). Pyramids and cascades: a synthesis of food chain functioning and stability. Ecology Letters, 22, 405–419.

Bezemer, T.M. & Jones, T.H. (1998). Plant-insect herbivore interactions in elevated atmospheric C02: quantitative analyses and guild effects. Oikos, 82, 212–222.

Binzer, A., Guill, C., Brose, U. & Rall, B.C. (2012). The dynamics of food chains under climate change and nutrient enrichment. Philosophical Transactions of the Royal Society B: Biological Sciences, 367, 2935–2944.

Binzer, A., Guill, C., Rall, B.C. & Brose, U. (2016). Interactive effects of warming, eutrophication and size structure: impacts on biodiversity and food-web structure. Global Change Biology, 22, 220–227.

Boit, A., Martinez, N.D., Williams, R.J. & Gaedke, U. (2012). Mechanistic theory and modelling of complex food-web dynamics in Lake Constance. Ecology Letters, 15, 594–602.

Borer, E.T., Bracken, M.E., Seabloom, E.W., Smith, J.E., Cebrian, J., Cleland, E.E. et al. (2013). Global biogeography of autotroph chemistry: is insolation a driving force? Oikos, 122, 1121–1130.

Brown, J.H., Gillooly, J.F., Allen, A.P., Savage, V.M. & West, G.B. (2004). Toward a metabolic theory of ecology. Ecology, 85, 1771–1789.

Cruz-Rivera, E. & Hay, M.E. (2000). Can quantity replace quality? Food choice, compensatory feeding, and fitness of marine mesograzers. Ecology, 81, 201–219.

Del Giorgio, P.A. & Gasol, J.M. (1995). Biomass Distribution in Freshwater Plankton Communities. The American Naturalist, 146, 135–152.

Diehl, S. (2007). Paradoxes of enrichment: effects of increased light versus nutrient supply on pelagic producer-grazer systems. The American Naturalist, 169, E173–E191.

Diehl, S., Berger, S. & Wöhrl, R. (2005). Flexible nutrient stoichiometry mediates environmental influences on phytoplankton and its resources. Ecology, 86, 2931–2945.

Droop, M. (1974). The nutrient status of algal cells in continuous culture. Journal of the Marine Biological Association of the United Kingdom, 54, 825–855.

Elser, J., Sterner, R., Gorokhova, E.a., Fagan, W., Markow, T., Cotner, J. et al. (2000). Biological stoichiometry from genes to ecosystems. Ecology Letters, 3, 540–550.

Elser, J.J., Bracken, M.E., Cleland, E.E., Gruner, D.S., Harpole, W.S., Hillebrand, H. et al. (2007). Global analysis of nitrogen and phosphorus limitation of primary producers in freshwater, marine and terrestrial ecosystems. Ecology Letters, 10, 1135–1142.

Elser, J.J., Loladze, I., Peace, A.L. & Kuang, Y. (2012). Lotka re-loaded: modeling trophic interactions under stoichiometric constraints. Ecological Modelling, 245, 3–11.

Elton, C. (1927). Animal ecology. Sidgwick & Jackson, LTD, London.

Englund, G., Ohlund, G., Hein, C.L. & Diehl, S. (2011). Temperature dependence of the functional response. Ecology Letters, 14, 914–921.

Enquist, B.J., West, G.B., Charnov, E.L. & Brown, J.H. (1999). Allometric scaling of production and life-history variation in vascular plants. Nature, 401, 907–911.

Falkowski, P.G., Barber, R.T. & Smetacek, V. (1998). Biogeochemical controls and feedbacks on ocean primary production. Science, 281, 200–206.

Finkel, Z.V., Beardall, J., Flynn, K.J., Quigg, A., Rees, T.A.V. & Raven, J.A. (2009). Phytoplankton in a changing world: cell size and elemental stoichiometry. Journal of Plankton Research, 32, 119–137.

Fussmann, K.E., Schwarzmüller, F., Brose, U., Jousset, A. & Rall, B.C. (2014). Ecological stability in response to warming. Nature Climate Change, 4, 206–210.

Gilarranz, L.J., Mora, C. & Bascompte, J. (2016). Anthropogenic effects are associated with a lower persistence of marine food webs. Nature communications, 7, 10737.

Gilbert, B., Tunney, T.D., McCann, K.S., DeLong, J.P., Vasseur, D.A., Savage, V. et al. (2014). A bioenergetic framework for the temperature dependence of trophic interactions. Ecology Letters, 17, 902–9014.

Hessen, D.O., Ågren, G.I., Anderson, T.R., Elser, J.J. & De Ruiter, P.C. (2004). Carbon sequestration in ecosystems: the role of stoichiometry. Ecology, 85, 1179–1192.

Hessen, D.O., Færøvig, P.J. & Andersen, T. (2002). Light, nutrients, and P:C ratios in algae: grazer performance related to food quality and quantity. Ecology, 83, 1886–1898.

Iles, A.C. (2014). Towards predicting community level effects of climate: relative temperature scaling of metabolic and ingestion rates. Ecology, 95, 2657–2668.

Irigoien, X., Huisman, J. & Harris, R.P. (2004). Global biodiversity patterns of marine phytoplankton and zooplankton. Nature, 429, 863–867.

Jensen, C.X. & Ginzburg, L.R. (2005). Paradoxes or theoretical failures? The jury is still out. Ecological Modelling, 188, 3–14.

Jeschke, J.M., Kopp, M. & Tollrian, R. (2004). Consumer-food systems: why Type I functional responses are exclusive to filter feeders. Biological Reviews, 79, 337–349.

Jonsson, T. (2017). Conditions for eltonian pyramids in Lotka-Volterra food chains. Scientific Reports, 7, 10912.

Kratina, P., Greig, H.S., Thompson, P.L., Carvalho-Pereira, T.S.A. & Shurin, J.B. (2012). Warming modifies trophic cascades and eutrophication in experimental freshwater communities. Ecology, 93, 1421–1430.

Lindeman, R.L. (1942). The trophic-dynamic aspect of ecology. Ecology, 23, 399–417.

Loladze, I., Kuang, Y. & Elser, J.J. (2000). Stoichiometry in producer-grazer systems: linking energy flow with element cycling. Bulletin of Mathematical Biology, 62, 1137–1162.

McAllister, C., LeBrasseur, R., Parsons, T. & Rosenzweig, M. (1972). Stability of enriched aquatic ecosystems. Science, 175, 562–565.

McCauley, D.J., Gellner, G., Martinez, N.D., Williams, R.J., Sandin, S.A., Micheli, F. et al. (2018). On the prevalence and dynamics of inverted trophic pyramids and otherwise top-heavy communities. Ecology Letters, 21, 439–454.

McCauley, E. & Kalff, J. (1981). Empirical relationships between phytoplankton and zooplankton biomass in lakes. Canadian Journal of Fisheries and Aquatic Sciences, 38, 458–463.

McCauley, E., Nisbet, R.M., Murdoch, W.W., de Roos, A.M. & Gurney, W.S.C. (1999). Large-amplitude cycles of Daphnia and its algal prey in enriched environments. Nature, 402, 653–656.

Montoya, J.M. & Raffaelli, D. (2010). Climate change, biotic interactions and ecosystem services. Philosophical Transactions of the Royal Society B: Biological Sciences, 365, 2013–2018.

Nelson, G.C. (2005). Millennium ecosystem assessment: drivers of ecosystem change: summary chapter. World Resources Institute, Washington, DC.

O’Connor, M.I., Piehler, M.F., Leech, D.M., Anton, A. & Bruno, J.F. (2009). Warming and resource availability shift food web structure and metabolism. PLoS Biology, 7, e1000178.

Petchey, O.L., McPhearson, P.T., Casey, T.M. & Morin, P.J. (1999). Environmental warming alters food-web structure and ecosystem function. Nature, 402, 69–72.

Peters, R.H. (1983). The ecological implications of body size. Cambridge University Press, Cambridge.

Rall, B.C., Brose, U., Hartvig, M., Kalinkat, G., Schwarzmüller, F., Vucic-Pestic, O. et al. (2012). Universal temperature and body-mass scaling of feeding rates. Philosophical Transactions of the Royal Society B: Biological Sciences, 367, 2923–2934.

Rastetter, E.B., Ågren, G.I. & Shaver, G.R. (1997). Responses of N-limited ecosystems to increased CO2: a balanced-nutrition, coupled-element-cycles model. Ecological Applications, 7, 444–460.

Rip, J.M.K. & McCann, K.S. (2011). Cross-ecosystem differences in stability and the principle of energy flux. Ecology Letters, 14, 733–740.

Robert W. Sterner, James J. Elser, Everett J. Fee, Stephanie J. Guildford & Thomas H. Chrzanowski (1997). The Light: Nutrient Ratio in Lakes: The Balance of Energy and Materials Affects Ecosystem Structure and Process. The American Naturalist, 150, 663–684.

Rosenzweig, M.L. (1971). Paradox of enrichment: destabilization of exploitation ecosystems in ecological time. Science, 171, 385–387.

Sarmiento, J.L., Slater, R., Barber, R., Bopp, L., Doney, S.C., Hirst, A. et al. (2004). Response of ocean ecosystems to climate warming. Global Biogeochemical Cycles, 18, 1–23.

Sentis, A., Binzer, A. & Boukal, D.S. (2017). Temperature-size responses alter food chain persistence across environmental gradients. Ecology Letters, 20, 852–862.

## References

Elton, C. (1927). Animal ecology. Sidgwick & Jackson, LTD, London.

Lindeman, R.L. (1942). The trophic-dynamic aspect of ecology. Ecology, 23, 399–417.

McCauley, E., Nisbet, R.M., Murdoch, W.W., de Roos, A.M. & Gurney, W.S.C. (1999). Large-amplitude cycles of *Daphnia* and its algal prey in enriched environments. Nature, 402, 653–656.

Menge, D.N.L., Hedin, L.O. & Pacala, S.W. (2012). Nitrogen and Phosphorus Limitation over Long-Term Ecosystem Development in Terrestrial Ecosystems. PLOS ONE, 7, e42045.

R Development Core Team (2017). R: a language and environment for statistical computing, R Foundation for Statistical Computing, Vienna, Austria.

Sentis, A., Hemptinne, J.L. & Brodeur, J. (2012). Using functional response modeling to investigate the effect of temperature on predator feeding rate and energetic efficiency. Oecologia, 169, 1117–1125.

Sentis, A., Hemptinne, J.L. & Brodeur, J. (2014). Towards a mechanistic understanding of temperature and enrichment effects on species interaction strength, omnivory and food-web structure. Ecology Letters, 17, 785–793.

Soetaert, K., Cash, J. & Mazzia, F. (2012). Solving differential equations in R. Springer Science & Business Media.

Sterner, R.W. & Elser, J.J. (2002). Ecological stoichiometry: the biology of elements from molecules to the biosphere. Princeton University Press.

Sterner, R.W. & Hessen, D.O. (1994). Algal nutrient limitation and the nutrition of aquatic herbivores. Annual Review of Ecology and Systematics, 25, 1–29.

Tabi, A., Petchey, O.L. & Pennekamp, F. (2019). Warming reduces the effects of enrichment on stability and functioning across levels of organisation in an aquatic microbial ecosystem. Ecology Letters, 22, 1061–1071.

Thomas, M.K., Aranguren-Gassis, M., Kremer, C.T., Gould, M.R., Anderson, K., Klausmeier, C.A. et al. (2017). Temperature–nutrient interactions exacerbate sensitivity to warming in phytoplankton. Global Change Biology, 23, 3269–3280.

Thomas, M.K., Kremer, C.T., Klausmeier, C.A. & Litchman, E. (2012). A global pattern of thermal adaptation in marine phytoplankton. Science, 338, 1085–1088.

Tilman, D. (1982). Resource competition and community structure. Princeton university press.

Tranvik, L.J., Downing, J.A., Cotner, J.B., Loiselle, S.A., Striegl, R.G., Ballatore, T.J. et al. (2009). Lakes and reservoirs as regulators of carbon cycling and climate. Limnology and Oceanography, 54, 2298–2314.

Tylianakis, J.M., Didham, R.K., Bascompte, J. & Wardle, D.A. (2008). Global change and species interactions in terrestrial ecosystems. Ecology Letters, 11, 1351–1363.

Uszko, W., Diehl, S., Englund, G. & Amarasekare, P. (2017). Effects of warming on predator–prey interactions – a resource-based approach and a theoretical synthesis. Ecology Letters, 20, 513–523.

Vasseur, D.A. & McCann, K.S. (2005). A mechanistic approach for modeling temperature-dependent consumer–resource dynamics. American Naturalist, 166, 184–198.

Vucic-Pestic, O., Ehnes, R.B., Rall, B.C. & Brose, U. (2011). Warming up the system: higher predator feeding rates but lower energetic efficiencies. Global Change Biology, 17, 1301–1310.

White, T. (1993). The inadequate environment. Nitrogen and the abundance of animals. Springer Verlag, Berlin.

Woods, H.A., Makino, W., Cotner, J.B., Hobbie, S.E., Harrison, J.F., Acharya, K. et al. (2003). Temperature and the chemical composition of poikilothermic organisms. Functional Ecology, 17, 237–245.

Yodzis, P. & Innes, S. (1992). Body size and consumer-resource dynamics. The American Naturalist, 139, 1151–1175.

Yvon-Durocher, G., Dossena, M., Trimmer, M., Woodward, G. & Allen, A.P. (2015). Temperature and the biogeography of algal stoichiometry. Global Ecology and Biogeography, 24, 562–570.

